# Peptidoglycan associated cyclic lipopeptide disrupts viral infectivity

**DOI:** 10.1101/635854

**Authors:** Bryan A. Johnson, Adam Hage, Birte Kalveram, Megan Mears, Jessica A. Plante, Sergio E. Rodriguez, Zhixia Ding, Xuemei Luo, Dennis Bente, Shelton S. Bradrick, Alexander N. Freiberg, Vsevolod Popov, Ricardo Rajsbaum, Shannan Rossi, William K. Russell, Vineet D. Menachery

**Affiliations:** Department of Microbiology and Immunology, University of Texas Medical Branch at Galveston; Department of Pathology, University of Texas Medical Branch at Galveston; World Reference Center for Emerging Viruses and Arboviruses, University of Texas Medical Branch at Galveston; Department of Biochemistry and Molecular Biology, University of Texas Medical Branch at Galveston; UTMB Electron Microscopy Laboratory, University of Texas Medical Branch at Galveston; UTMB Mass Spectrometry Facility, University of Texas Medical Branch at Galveston; Institute for Human Infections and Immunity, University of Texas Medical Branch at Galveston

## Abstract

Enteric viruses exploit bacterial components including lipopolysaccharides (LPS) and peptidoglycan (PG) to facilitate infection in humans. With origins in the bat enteric system, we wondered if severe acute respiratory syndrome-coronavirus (SARS-CoV) or Middle East respiratory syndrome-CoV (MERS-CoV) also use bacterial components to modulate infectivity. To test this question, we incubated CoVs with LPS and PG and evaluated infectivity finding no change following LPS treatment. However, PG from *B. subtilis* reduced infection >10,000-fold while PG from other bacterial species failed to recapitulate this. Treatment with an alcohol solvent transferred inhibitory activity to the wash and mass spectrometry revealed surfactin, a cyclic lipopeptide antibiotic, as the inhibitory compound. This antibiotic had robust dose- and temperature-dependent inhibition of CoV infectivity. Mechanistic studies indicated that surfactin disrupts CoV virion integrity and surfactin treatment of the virus inoculum ablated infection *in vivo*. Finally, similar cyclic lipopeptides had no effect on CoV infectivity and the inhibitory effect of surfactin extended broadly to enveloped viruses including influenza, Ebola, Zika, Nipah, Chikungunya, Una, Mayaro, Dugbe, and Crimean-Congo hemorrhagic fever viruses. Overall, our results indicate that peptidoglycan-associated surfactin has broad virucidal activity and suggest bacteria byproducts may negatively modulate virus infection.

**Importance:** In this manuscript, we considered a role for bacteria in shaping coronavirus infection. Taking cues from studies of enteric viruses, we initially investigated how bacterial surface components might improve CoV infection. Instead, we found that peptidoglycan-associated surfactin is a potent viricidal compound that disrupts virion integrity with broad activity against enveloped viruses. Our results indicate that interactions with commensal bacterial may improve or disrupt viral infections highlighting the importance of understanding these microbial interactions and their implications for viral pathogenesis and treatment.

## Introduction

Commensal bacteria inhabit nearly every surface of the human body, influencing numerous host processes (1, 2). While considered to serve a protective role, recent studies indicate enteric viruses exploit bacterial envelope components to facilitate infection (3). Poliovirus was found to bind both lipopolysaccharides (LPS) and peptidoglycan (PG) to enhance its thermostability and receptor affinity, facilitating *in vivo* infection (3). Antibiotic depletion of commensal bacteria inhibited oral poliovirus infection, but was rescued by recolonization, pretreatment of virus with LPS, or bypassing the enteric system through intraperitoneal injection (3). Other viruses including reovirus, mouse mammary tumor virus, and murine norovirus have been shown to use similar mechanisms to facilitate infection (3, 4). Together, these results indicate a key role for commensal bacteria in improving infectivity and pathogenesis of enteric viruses.

Like the enteric system, the respiratory tract harbors high levels of commensal bacteria (1). Given the origins of Severe Acute Respiratory Syndrome-Coronavirus (SARS-CoV) and Middle East Respiratory Syndrome (MERS)-CoV in the bat enteric system (5), we wondered if CoVs utilized bacterial components to facilitate infection. Previous work had identified a key role for the TLR pathways in immunity to SARS-CoV with the absence of LPS binding TLR4 or its downstream adaptors resulting in augmented disease (6–8). Given the interactions observed between enteric viruses and bacterial components, CoVs may also use similar microbial components to improve infectivity and subsequently stimulate the TLR4 response.

In this study, we explored the relationship between bacterial surface components and CoV infection. Surprisingly, we found PG from *Bacillus subtilis* reduced CoV infectivity. Using mass spectrometry, we identified a cyclic lipopeptide (CLP), surfactin, as the molecule responsible for CoV inhibition. Surfactin’s inhibitory effect was dose and temperature dependent with treatment disrupting the integrity of the CoV particle. Notably, surfactin treatment of the inoculum ablated CoV infection *in vivo*, but prophylactic treatment had no effect. Other similar CLPs had no effect on CoV infectivity suggesting surfactin’s virucidal properties were unique. Importantly, surfactin treatment reduced the infectivity of several other enveloped viruses, including influenza A, Zika, Dugbe, Nipah, Crimean-Congo Hemorrhagic Fever, chikungunya, Mayaro, Una, and Ebola viruses. Together, these results demonstrate the efficacy of surfactin as a virucidal compound and highlight the potential for microbial environment to modulate virus infection.

## Results

### Peptidoglycan derived from *B.* subtilis reduces with coronavirus infectivity

Given their origins in bat enteric systems, we wondered if CoVs might be stabilized by bacterial components (5). To test this possibility, Human CoV-229E, a common cold associated CoV, and MERS-CoV were treated with control (PBS), LPS (*Escherichia coli)* or PG (*Bacillus subtilis)* and viral infectivity was determined (Fig. 1*A*). In contrast to enteric viruses, LPS had no effect on CoV infectivity; however, the presence of PG from *B. subtilis* dramatically reduced the infectivity of both HCoV-229E and MERS-CoV (Fig. 1*B*). The structure of PG varies considerably between bacterial species (9), suggesting that PG from different bacteria may have distinct effects on CoV infectivity. To explore this, we tested a diverse set of bacterial derived PGs for the ability to modulate CoV infection (Fig. 1*C*). Notably, only PG derived from *B. subtilis* reduced HCoV-229E and MERS-CoV infection, suggesting that interference with CoV infectivity is not shared by PG from all bacterial species.

**Figure 1:**
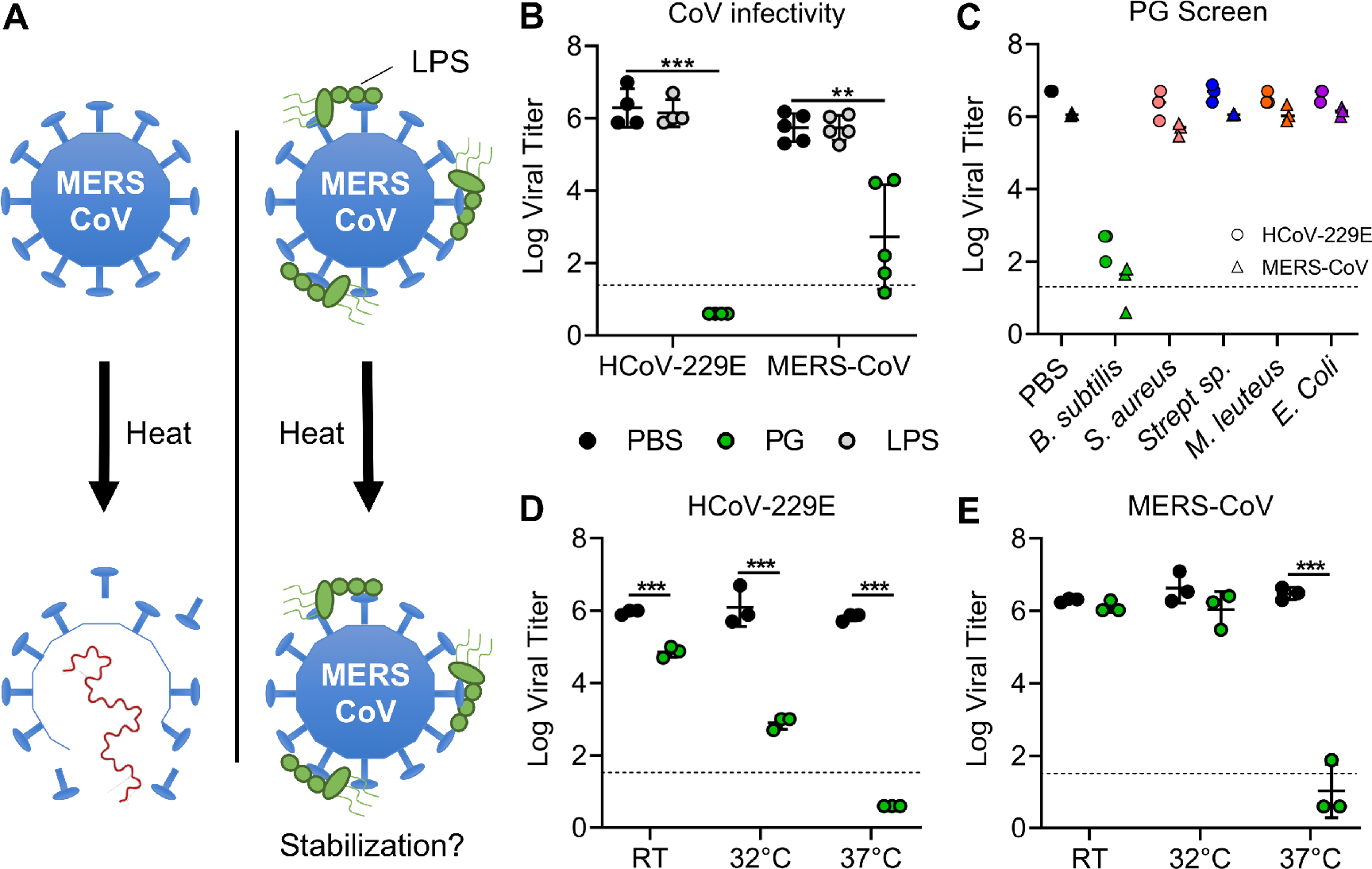
Peptidoglycan from *Bacillus subtilis* reduces coronavirus infectivity. (A) Bacterial envelope components such as LPS are bound to CoVs, increasing their thermostability (right) relative to untreated samples (left). (B) Relative infectivity of HCoV-229E (n=4) and MERS-CoV (n=5) after treatment with PBS alone (black), LPS from *Escherichia coli* (grey), or PG from *B. subtilis* (green) following 2 hour incubation at 37°C. (C) HCoV-299E (circles) and MERS-CoV (triangles) infectivity after treatment for 2 hours at 37°C with peptidoglycan from the indicated bacterial species (n=3). (D) HCoV-229E and (E) MERS-CoV after treatment with PG from *B. subtilis* at room temperature (RT), 32°C, and 37°C (n=3). For all dot plots, the centered bar represents the group mean while the error bars represent SD. P-values are based on the two-tailed Student’s *t* test as indicated: * *P* < 0.05, ** *P* < 0.01, *** *P* < 0.001.

Next, we wondered if incubation temperature also played a role in *B. subtilis* PG reduction of CoV infectivity. To investigate, HCoV-229E and MERS-CoV stocks were treated with *B. subtilis* PG at room temperature (RT), 32°C, or 37°C (Fig. 1*D-E*). Interestingly, PG disruption of viral infectivity was reduced at lower temperatures. For HCoV-229E, infectivity had a step-wise reduction with increasing temperature (Fig. 1*D*). In contrast, PG reduction of MERS-CoV infectivity was ablated at lower temperatures, with no significant loss of viral infectivity at either RT or 32°C (Fig. 1*E*). Together, these data indicate that the inhibitory effect of *B. subtilis* PG is influenced by incubation temperature.

### Infectivity inhibition can be disassociated from PG

Two possible scenarios explain why only *B. subtilis* PG reduces CoV infectivity: 1) *B. subtilis* PG reduces infectivity directly, using unique structural features absent in PG from other bacteria; or 2) the PG preparation contains another compound that mediates inhibition. To differentiate these possibilities, we exploited the poor solubility of PG, washing it in a variety of solvents to separate its inhibitory effect (Fig. 2A). After three washes in PBS, PG maintained its reduction of HCoV-229E infectivity (Fig. 2*A*). In contrast, PG washed with either 100% ethanol or DMSO lost the ability to inhibit HCoV-229E infectivity (Fig. 2*A*). These results suggest that the washes either modified the inhibitory capacity of PG or removed a soluble compound responsible for reducing CoV infectivity. To explore this, the supernatants from clarified PG samples were incubated with HCoV-229E (Fig. 2*B*). While the PBS, PBS control, and ethanol control had no inhibitory effect, the ethanol supernatant from PG potently reduced viral infectivity of HCoV-229E (Fig. 2*B*). Together, these data indicate that a soluble compound distinct from, but present in the PG sample, is responsible for reducing CoV infectivity.

**Figure 2:**
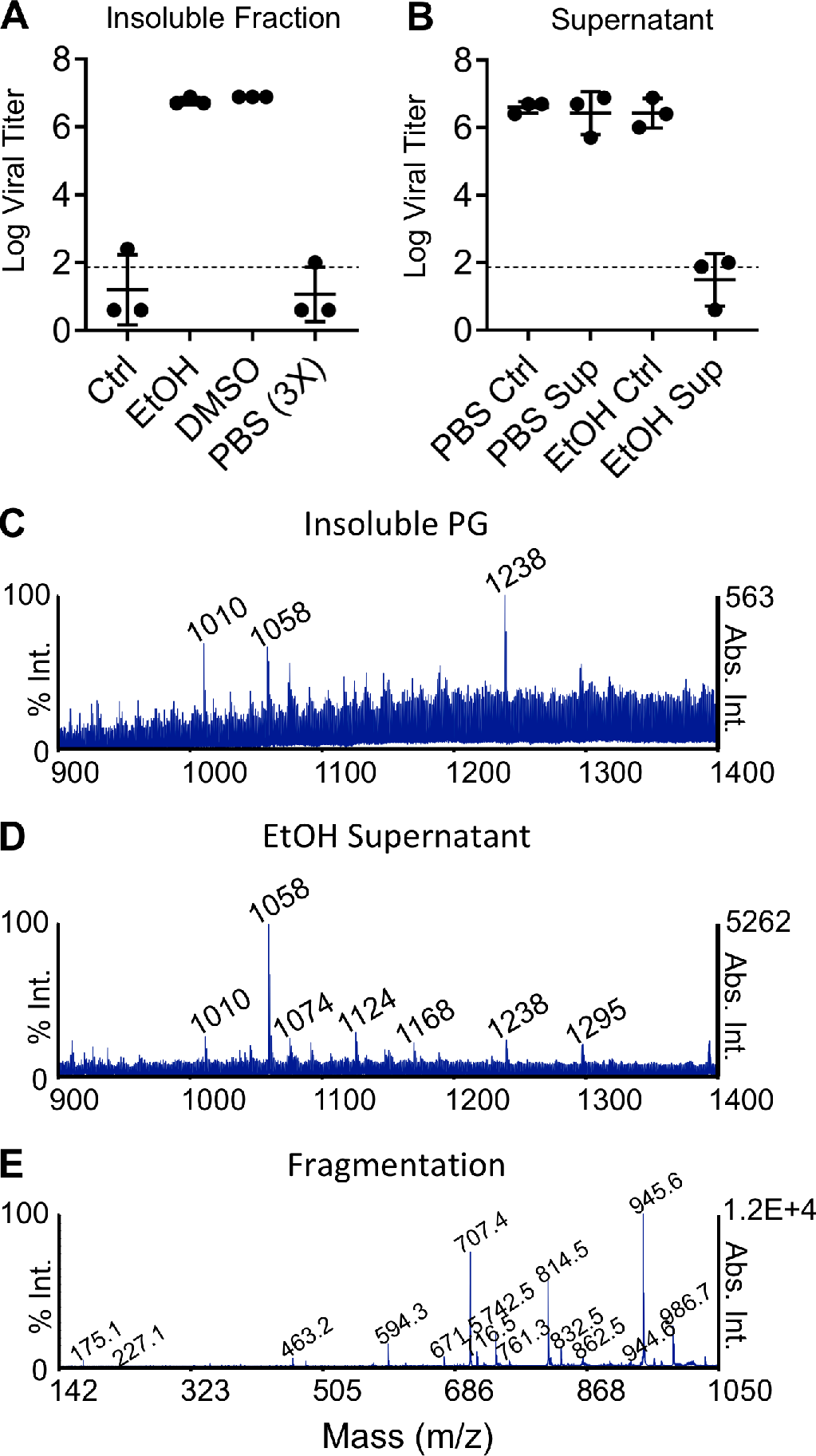
Identification of surfactin from *B. subtilis* peptidoglycan. (A and B) PG from *B. subtilis* in a PBS solution was clarified, washed with the indicated solvents, and clarified again. Supernatants were decanted and retained, while the insoluble fractions were resuspended in PBS. The (A) insoluble fraction and (B) supernatants were then used to treat HCoV-229E and relative infectivity determined (n=3). (C, D, and E) Mass spectrometry performed was performed on PG (C) and ethanol wash (D). The peak corresponding to the molecular mass 1058 in the ethanol wash was then further fragmented (E) to determine the identity of the molecule. Representative spectra shown. For all dot plots, the centered bar represents the group mean and error bars the SD.

### Mass spectrometry identifies the inhibitor as surfactin

Having isolated the inhibitory molecule, we utilized mass spectrometry to determine its identity. Unwashed *B. subtilis* PG and ethanol supernatants were analyzed using MALDI-TOF/TOF mass spectrometry (Fig. 2*C*). In the PG samples, three prominent peaks were observed with masses of 1010.5, 1058.7, and 1238.6 (Fig. 2*C*). While all of these peaks were present in the ethanol supernatant, the compound with a mass 1058.7 was enriched nearly 10-fold (Fig. 2*D*). Further analysis of this peak by fragmentation produced a spectrum matching that of the cyclic lipopeptide surfactin (10) a potent biosurfactant produced naturally by *B. subtilis* and shown previously to have antimicrobial and antiviral properties (11, 12) (Fig 2*E*, for structure see Fig 6*A*). Given its abundance, enrichment in the ethanol wash, as well as its described antiviral properties, we concluded that surfactin likely conferred the *B. subtilis* PG with the ability to interfere with CoV infection.

### Reduction of CoV infectivity by surfactin is temperature” and dose-dependent

To confirm its inhibitory effect, we characterized the ability of purified surfactin to reduce CoV infectivity. HCoV-229E, MERS-CoV, or SARS-CoV were treated with PBS or surfactin at either RT, 32°C, or 37°C. For all three CoVs, surfactin reduced infectivity after treatment at 37°C, with a near complete loss of infectious virus (Fig. 3*A-C*). Similar to *B. subtilis* PG, the degree of reduction varied based on incubation temperature and varied between the CoVs (Fig. 3*A-C*). To further characterized the kinetics of inhibition, HCoV-229E and MERS-CoV were treated with surfactin at 4°C, RT, 32°C, or 37°C and sampled over a time-course (Fig. 3*D-E*). At both 32°C and 37°C, surfactin rapidly reduced HCoV-229E and MERS-CoV, with a near complete loss of infectivity after two hours of treatment (Fig. 3*D-E*). In contrast RT incubation reduced CoV infectivity more slowly and, surfactin’s effects were ablated at 4°C. We also observed dose dependent changes in surfactin activity against HCoV-229E, MERS-CoV, and SARS-CoV (Fig. 3*F*). Interestingly, higher concentrations of surfactin were required for inhibition of HCoV-229E when compared to either SARS-CoV or MERS-CoV, whose inhibition curves were nearly identical. Together, these data indicate that both temperature and dose impact surfactin’s inhibitory effects.

**Figure 3:**
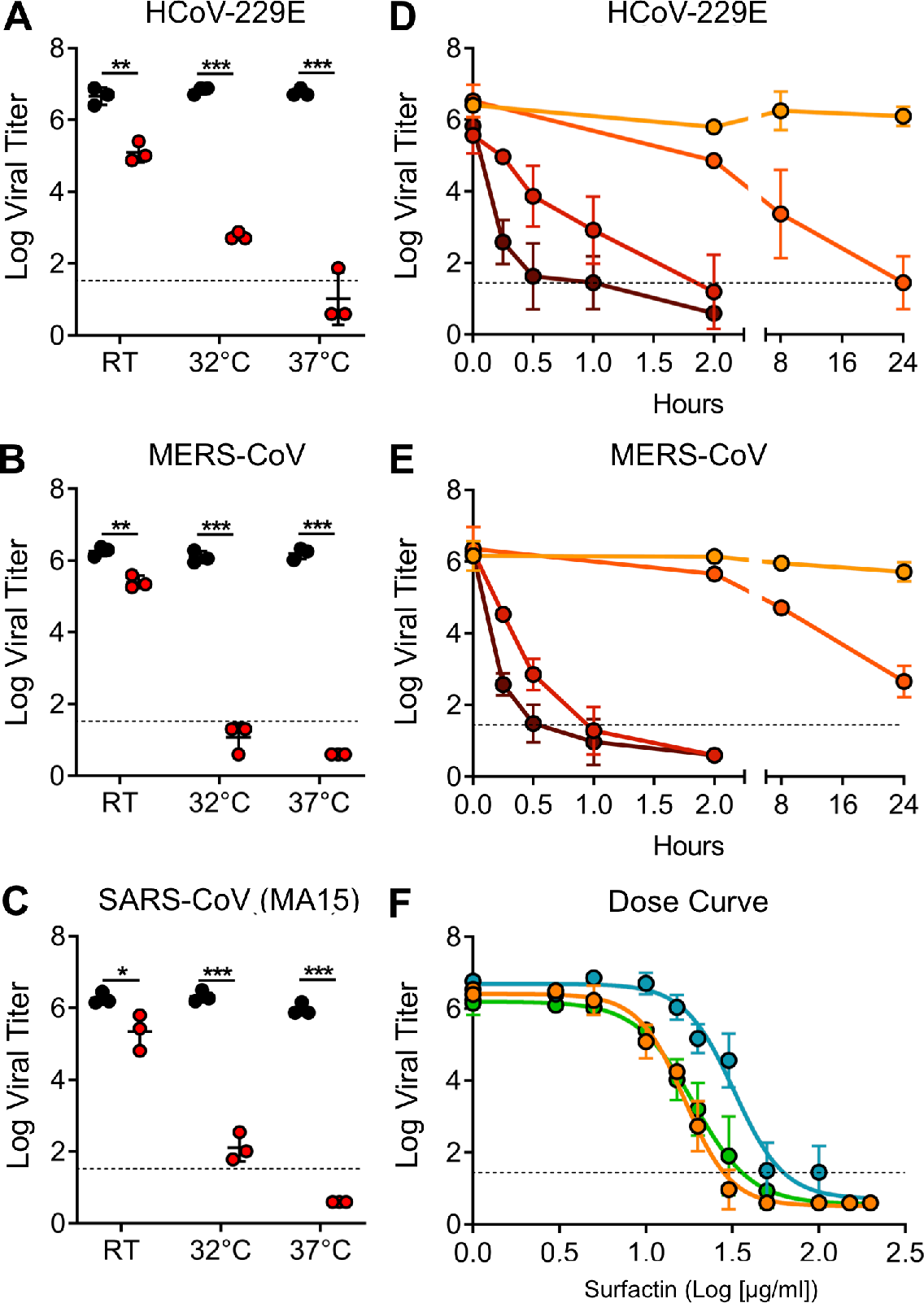
Characterization of CoV inhibition by surfactin. (A) HCoV-229E, (B) MERS-CoV, and (C) SARS-CoV (MA15) were treated PBS alone (black) or surfactin (red) and at room temperature (RT), 32°C, and 37°C and infectivity determined (n=3). HCoV-229E (D) and MERS-CoV (E) were treated for the indicated time at 4°C (light orange), RT (dark orange), 32°C (light red), or 37°C (dark red) and infectivity determined (n=3). (F) HCoV-229E (blue), MERS-CoV (orange), and SARS-CoV MA15 (green) were diluted over a range of concentrations of surfactin and viral infectivity determined (n=3). For dot plots, each point represents the titer from an independent experiment while the group mean is indicated by a line. Each point on the line graph represents the group mean. All error bars represent SD. The two tailed students t-test was used to determine P-values: * *P* < 0.05, ** *P* < 0.01, *** *P* < 0.001.

### Surfactin reduces CoV infectivity by disrupting the structural integrity of viral particles

Prior studies with surfactin offers two mechanisms for virucidal activity: disruption of the viral membrane or inhibition of host-virus membrane fusion (11–13). To determine if virion integrity was maintained, we performed RNase I protection assays. Following surfactin treatment, particles were exposed to RNase I to digest exposed viral RNA; samples were subsequently extracted for RNA and relative viral RNA determined by quantitative reverse transcription real time PCR (RT-qPCR). Increasing surfactin concentrations correlated with a decrease in viral RNA and viral titer for both HCoV-229E (Fig. 4*A*) and MERS-CoV (Fig. 4*B*). These results indicate that disruption of virion integrity is the primary mechanism by which surfactin inhibits CoV infection. To confirm these results, we performed transmission electron microscopy (TEM) on HCoV-229E treated with surfactin or PBS. In PBS-treated samples, numerous intact HCoV-229E particles could be visualized (Fig. *4C*-*D*); in contrast, no viral particles were found in any of the surfactin treated samples (Fig. *4D*). Taken together these results demonstrate that surfactin inhibits CoV infection primarily through the disruption of viral particles.

**Figure 4:**
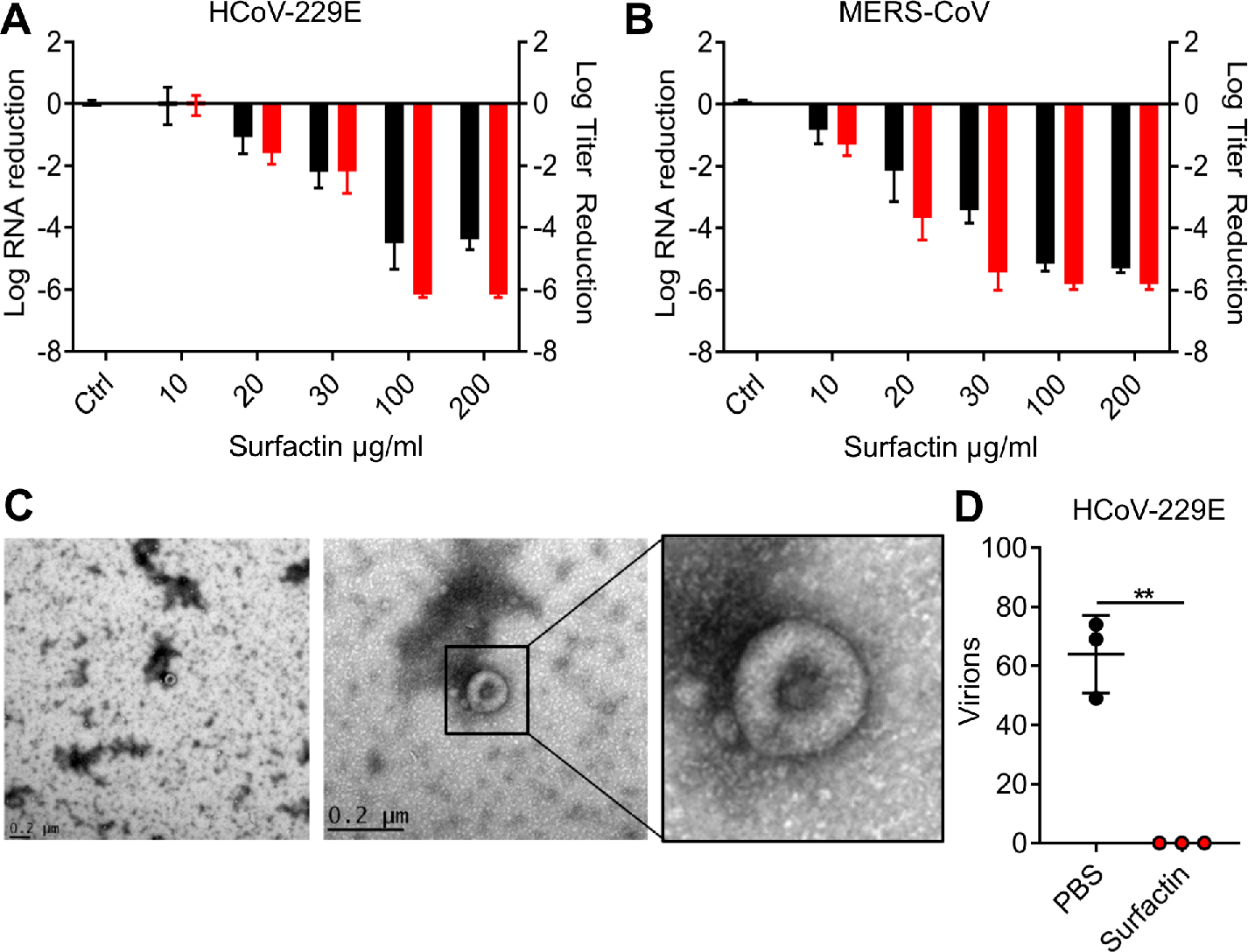
Surfactin disrupts CoV structural integrity. (A) HCoV-229E and (B) MERS-CoV were treated with the indicated concentrations of surfactin. Viral infectivity was then determined (red) or samples were then treated with RNAse I, RNA extracted, and viral genome copy number determined by RT-qPCR (black). (C and D) PBS or surfactin treated HCoV-229E samples negatively stained and examined by TEM and virions counted (n=3). Representative micrograph shown in (C) while total counts are displayed in (D). Horizontal lines represent group mean while error bars represent SD. Two-tailed students t-test determined significance: * *P* < 0.05, ** *P* < 0.01, *** *P* < 0.001.

### *In vivo* characterization of surfactin on CoV infection

With no approved therapeutics (14), emerging, zoonotic CoVs pose a significant threat to public health (15, 16). Therefore, we wanted to examine the potential of surfactin to treat infections *in vivo*. We tested whether direct treatment of the inoculum reduced *in vivo* infection and disease. SARS-CoV (10^4^ plaque forming units (PFU)) was treated with PBS or surfactin and used to infect BALB/c mice intranasally (IN). Mice were monitored over 4 days for weight loss and lethality, with lung titers determined at 2- and 4-days post infection. As expected, animals infected with PBS treated virus experienced rapid weight loss and exhibited high lung titers at both 2- and 4-days post infection (Fig. 5A). In contrast, mice infected with surfactin treated SARS-CoV lost no weight and no infectious virus was detected in their lungs (Fig. 5*B*). Additionally, mock infected mice receiving surfactin alone demonstrated no signs of disease or weight loss, suggesting that surfactin treatment alone does not have any pathological effects (Fig. 5*A*).

**Figure 5:**
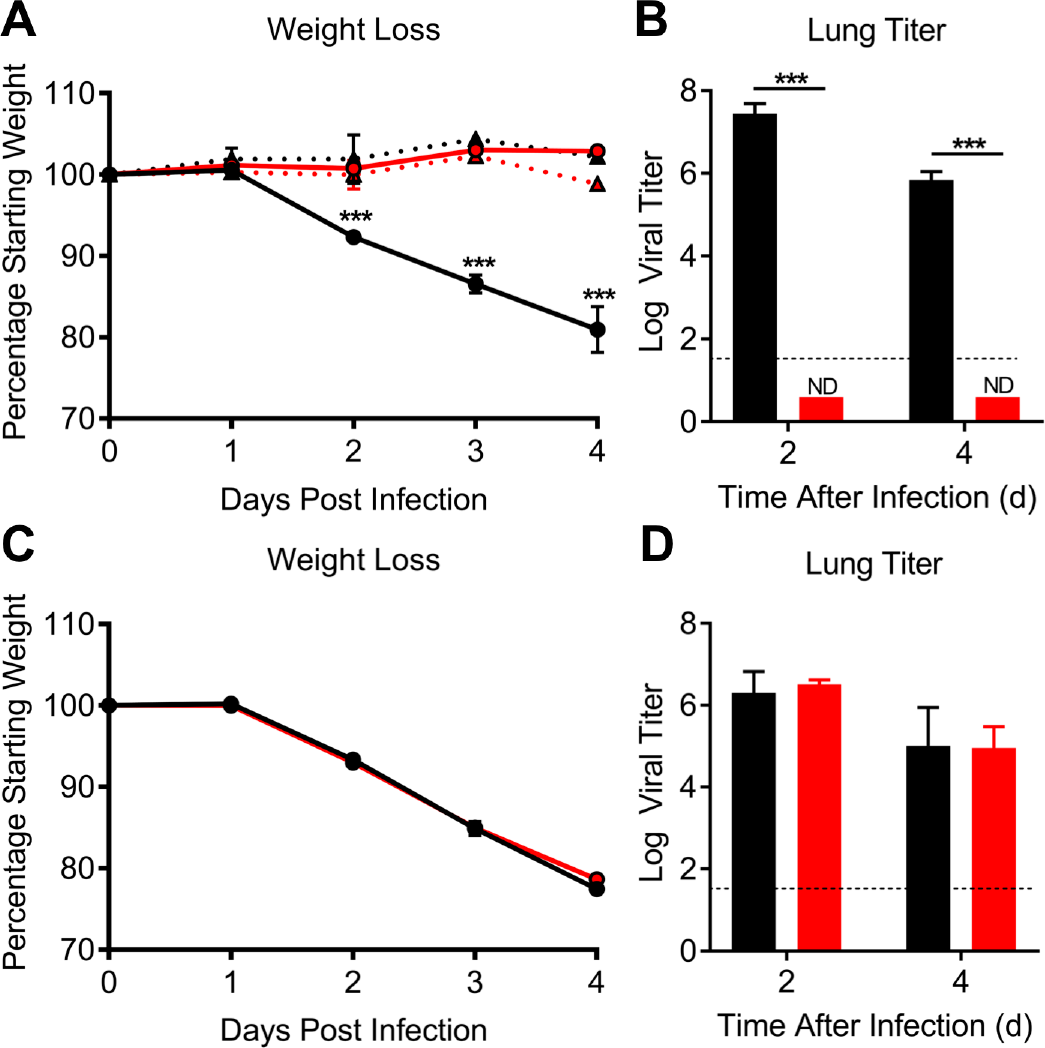
*In vivo* characterization of surfactin treatment on SARS-CoV infection. **(A and B)** BALB/C mice were then intranasally infected with 10^4^ PFU of PBS (black) or surfactin (red) treated SARS-CoV MA15 and (A) monitored for weight loss over 4 days. Dotted lines and triangles represent mock infected animals with PBS alone (black) or surfactin alone (red). (B) Lung tissue was harvested and viral titer determined at day 2 and day 4. n=4 for all infected groups, n=2 for mock groups. (C and D) BALB/C mice were pretreated intranasally with 50 μl of either PBS (black) or surfactin in PBS (red). 18 hours later, BALB/C mice were infected with 10^4^ PFU of SARS-CoV (MA15) and (C) monitored for weight loss over 4 days. (D) Lung titer determined 2-(n=5) and 4-days post infection (n=10). Dots on line graphs and bars on bar graphs represent the group mean. ND indicates that no titers were detected. All error bars represent SD. P-values were calculated using the two-tailed student’s t-test, with: * *P* < 0.05, ** *P* < 0.01, *** *P* < 0.001.

To examine therapeutic potential, we next evaluated if pretreatment with surfactin could reduce respiratory CoV disease. BALB/c mice were treated IN with 50 μl of either PBS control or surfactin daily, starting 18 hours prior to infection, and continuing over the first two days of infection. Mice were subsequently infected with 10^4^ PFU of SARS-CoV (MA15) and monitored for weight loss and lethality, with lung titer determined at 2 and 4 days post-infection. In contrast to the surfactin-treated inoculum, prophylactic surfactin treatment had no effect on weight loss (Fig. 5*C*) or viral titer in the lung (Fig. 5*D*). These results indicate that prophylactic surfactin treatment by this route does not reduce SARS-CoV disease in this mouse model.

### Effects of other cyclic lipopeptides on CoV infectivity

Surfactin belongs to a family of 80 natural antibiotic compounds referred to as cyclic lipopeptides (CLPs) (11). While structurally diverse, all CLPs share two key features: a non-polar hydrocarbon tail and a non-ribosomally produced peptide ring (11, 12). While many CLPs have been found to be antifungal and antibacterial, antiviral properties have not been described except for surfactin, (11, 12). Therefore, we tested six CLPs for the ability to reduce CoV infectivity (Fig. 6*A*). Despite similar biochemical structures, none of the CLPs tested had a significant effect on HCoV-229 or MERS-CoV infection (Fig. 6B). These results suggest that unique features allow surfactin to reduce CoV infectivity.

**Figure 6:**
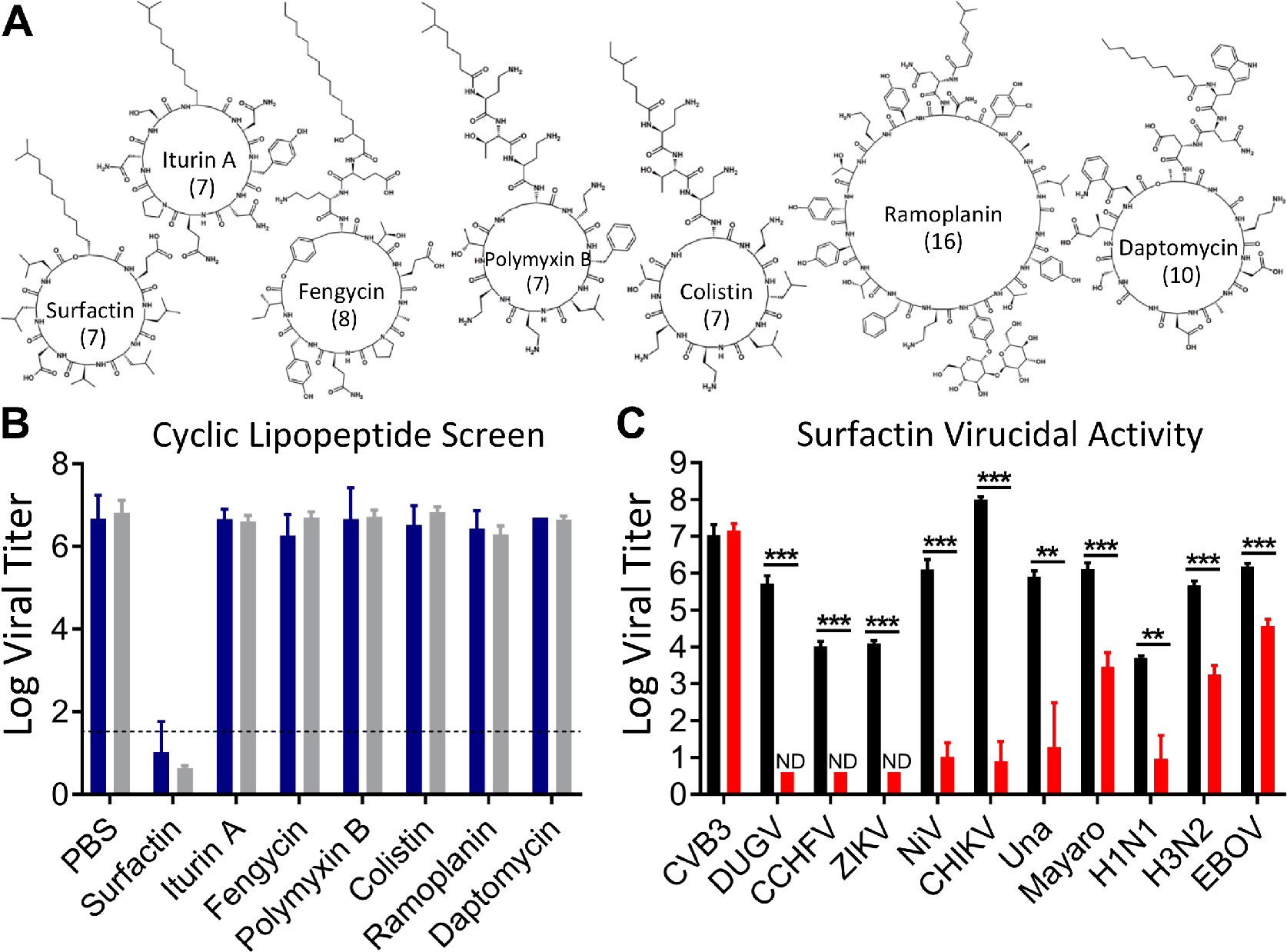
Surfactin, but not other cyclic lipopeptides, broadly reduce the infectivity of enveloped viruses. (A) Biochemical models of each of the seven cyclic lipopeptides tested. The number of amino acids present in the cyclic ring are shown in parentheses. (B) HCoV-229E (blue) and MERS-CoV (grey) were treated with PBS or the indicated cyclic lipopeptides in PBS and incubated for 2 hours at 37°C. Viral infectivity was then determined (n=3). (C) The indicated viruses were diluted PBS (black) or surfactin (red), incubated for 2 hours at 37°C, and viral infectivity determined (n=3). Viruses are abbreviated as follows: Coxsackievirus (CVB3), Dugbe (DUGV), Crimean-Congo hemorrhagic fever (CCHFV), Zika (ZIKV), Nipah (NiV), and Chikungunya (CHIKV), Una, Mayaro, Influenza A strains H1N1 and H3N2, and Ebola (EBOV). Bar graph bars represent the group means. Error bars represent SD. ND indicates that no titers were detected. The student’s t-test was used to calculate p-values, with: * *P* < 0.05, ** *P* < 0.01, *** *P* < 0.001.

### Surfactin broadly reduces viral infectivity

With its potent antiviral properties against CoVs, we wanted to test surfactin’s effect against other highly pathogenic viruses. Given its ability to disrupt virion integrity, we focused on enveloped viruses from diverse families including two Influenza A strains (H1N1, H3N2), Zika Virus (ZIKV), Dugbe Virus (DUGV), Nipah Virus (NiV), Crimean-Congo Hemorrhagic Fever Virus (CCHFV), Chikungunya Virus (CHIKV), Mayaro virus, Una virus, and Ebola virus (EBOV). As a negative control, we tested the non-enveloped Coxsackievirus B3 (CVB3). Each virus was treated with either PBS or surfactin and viral infectivity determined. As expected, surfactin had no effect on the non-enveloped CVB3 (Fig. 6*C*). In contrast, surfactin significantly reduced infectivity in each of the enveloped viruses (Fig. 6*C*), but the magnitude of effect was not uniform. Most enveloped viruses were reduced either below their limit of detection or greater than 100,000-fold. In contrast, Mayaro, both influenza strains, and EBOV exhibited some resistance, having their infectious titer reduced only 2.6, 2.7, 2.4, and 1.6 logs, respectively. These data suggest that while surfactin treatment broadly reduced the infectivity of enveloped viruses, factors beyond the mere presence or absence of an envelope may govern overall sensitivity.

## Discussion

In this study, we explored the relationship between bacterial components and CoV infection. While initially predicting enhanced infection, treatment with *B. subtilis* PG reduced CoV infectivity, while envelope components from other bacteria had no effect. Separating the inhibitory effect using solvent washes, we used mass spectrometry to identify that the CLP surfactin was responsible for reduced CoV infectivity and disruption of virion integrity. Unfortunately, despite efficacy against the inoculum, prophylactic surfactin treatment prior to infection had no effect on CoV related disease *in vivo*. Notably, other CLPs had no effect on CoV infectivity despite having similar biochemical structures. Finally, we found that surfactin treatment was efficacious against many enveloped viruses *in vitro* including IAV strains H1N1 and H3N2, ZIKV, DUGV, NiV, CCHFV, CHIKV, Una, Mayaro, and EBOV. Together, these data demonstrate that surfactin is a potent virucide and highlight that interactions with bacterial derived compounds can also negatively modulate virus infection.

Over the last two decades, surfactin has been shown to be anti-bacterial, anti-fungal, and anti-viral (11, 12, 17–19). Mechanistically, surfactin’s broad anti-microbial efficacy has been linked to disruption of lipid membranes (13). However, more recently, researchers described surfactin’s efficacy against the animal CoV porcine epidemic diarrhea virus (PEDV), and suggested that surfactin inhibited viral-host membrane fusion (19). In contrast to the PEDV results, we found that surfactin treatment disrupted virion integrity, exposing the viral RNA to RNase I mediated degradation (Fig. 4*A-B*). Transmission electron microscopy (TEM) confirmed the absence of intact virions in surfactin treated samples (Fig. 4*C*). Thus, while both PEDV and human CoVs are sensitive to surfactin treatment, infectivity reduction may be the result of different mechanisms due to differences the surfactin dose, virion composition, tissue environment (respiratory vs enteric), or other factor.

Similar to the question of mechanism, *in vivo* efficacy of surfactin also varied between PEDV and human CoVs. While surfactin ablates SARS-CoV disease when treating the inoculum, prophylactic treatment was not protective. Why surfactin failed to protect mice against SARS-CoV is puzzling, given that efficacy of prophylactic oral surfactin treatment against PEDV disease (19). One explanation is the physical environment of the respiratory and gastrointestinal tracts differs significantly, making intranasal (IN) surfactin administration ineffective due to tissue specific differences. Thus, while oral surfactin administration may be effective at delivering surfactin to infected gastrointestinal tissue, IN administration may not be as effective, particularly in the lower parts of the lung. Alternative delivery methods such as the inhalation of an aerosolized surfactin may overcome these problems. Additionally, several surfactin derivatives exist and may enhance its virucidal activity in the context of respiratory infection *in vivo* (11, 12).

In addition to CoVs, we examined surfactin’s virucidal efficacy against other enveloped viruses, discovering broad efficacy, but wide variation. While all tested enveloped viruses were sensitive to surfactin treatment, IAV strains H1N1 and H3N2, Mayaro, and EBOV demonstrated a degree of resistance. These data suggest that factors beyond the mere presence of a viral envelope regulate surfactin efficacy. One possible factor is the lipid content of the viral envelope. Previous studies have shown that membranes enriched in cholesterol and phosphatidylethanolamines (PE) are resistant to surfactin permeabilization, while membranes containing phosphatidylcholines (PC) are more sensitive (20). The envelope of Influenza A viruses have been reported to be enriched for both cholesterol and PE (21), providing support for this hypothesis. Unfortunately, the lipid content of the other viruses tested have not been determined, preventing direct comparison. Nevertheless, some broad observations are worth mentioning. CoVs, ZIKV, and bunyaviruses (CCHFV and DUGV) derive their envelopes from either the Golgi Apparatus and Endoplasmic Reticulum, organelles enriched in surfactin sensitive PC (22). NiV (23), IAV (24), EBOV (25) are thought to derive their envelopes from lipid rafts of the plasma membrane, which could specify their lipid content and thus surfactin sensitivity. Alphaviruses such as CHIKV, Mayaro, and Una also bud form the plasma membrane, though neither the lipid content, nor the involvement of lipid rafts has been explored (26). Together, these observations suggest the lipid content of enveloped viruses may explain their differential sensitivity to surfactin.

The failure of other CLPs to reduce CoV infectivity is also surprising, given the structural similarity to surfactin. In particular, Iturin A is both biochemically similar to surfactin and has also been reported to disrupt lipid membranes (Fig. 6*A*) (11). A possible explanation involves differences in their mechanisms of action. Surfactin penetrates lipid layers, alone solubilizing and permeabilizing them (13). In contrast, Iturin A must interact with sterol components to cause membrane permeabilization, explaining its broad anti-fungal, but only selective antibacterial activity (11). However, Iturin A is also quite hemolytic (11, 12), making it unclear why the membranes of enveloped viruses grown in mammalian cells would not also be susceptible to this mechanism, due to the presence of sterols. Compounding this mechanistic uncertainty, daptomycin’s permeabilization of membranes requires no such interaction, but CoVs are resistant to its effects as well (27) (Fig. 6*B*). The results argue that surfactin possesses unique properties conferring its virucidal activity. Surfactin is known to adopt a unique β-sheet like “horse-saddle” conformation, which may facilitate membrane permeabilization (13). Molecular dynamics simulations suggest temperature directly regulates the openness of the horse saddle structure and may explain why surfactin’s virucidal activity is also temperature sensitive (28). In total, these results highlight our poor understanding of membrane disruption by CLPs and argues that biochemical studies of these compounds inhibition of enveloped viruses are needed.

While the microbiome has historically been thought to serve a protective role against pathogens (1, 2), recent studies with viruses complicate this view. Studies with poliovirus demonstrated that the presence of commensal bacteria is necessary for oral poliovirus infection in mice (29). Similar findings with other enteric viruses suggest that utilizing bacterial components is a common approach. In contrast, our results add further complexity, demonstrating that surfactin, a secondary metabolite of *B. subtilis*, can potently reduce CoV infectivity. Though *B. subtilis* is not generally part of the human microbiome (30), it is often used as an intestinal probiotic and has been found to transiently persist in the gut (31). Additionally, surfactin-like molecules are produced by a broad array of bacterial species (11, 32–34). For example, the novel surfactin like CLP Coryxin was recently found to be produced by *Corynebacterium xerosis*, a common member of the respiratory microbiome (34). These facts suggest that microbial components typically thought to work against bacterial competitors could also potentially disrupt viral infection. Thus, as the relationship between the microbiome and viral infections is further explored, the role bacterial metabolites such as surfactin and other CLPs play in modulating infection must be considered in viral disease. Overall, these results highlight the dynamic microbial environment and its potential to impact viral pathogenesis as well as identify novel inhibitory factors for therapeutic use.

## Materials and Methods

### Viruses, cells, and *in vitro* infection

HCoV-229E, provided by the World Reference Center for Emerging Viruses and Arboviruses (WRCEVA), was propagated on HUH7 cells grown in DMEM (Gibco), 10% fetal bovine serum (Hyclone), and 1% antibiotic-antimycotic (A/A) (Gibco). Titration was performed by TCID_50_ in HUH7 cells and calculated by the Spearman-Karber Method. MERS-CoV (EMC-2012 strain) (35) and recombinant SARS-CoV (MA15) (36) were titrated and propagated on VeroCCL81 and VeroE6 cells, respectively, grown in DMEM with 5% fetal bovine serum and 1% A/A. Standard plaque assays were used for SARS- and MERS-CoV (37, 38). Cocksackievirus B3 (39), chikungunya (40), Nipah (41), Dugbe (42), Zika (43), Crimean-Congo hemorrhagic fever (42), Influenza A H1N1(A/California/04/09) H3N2 (A/Panama/2007/99) (44), and Ebola viruses (45) were propagated and quantitated via standard methods. All experiments involving infectious virus were conducted at the University of Texas Medical Branch (Galveston, TX) in approved biosafety level 2, 3, or 4 (BSL) laboratories and animal facilities, with routine medical monitoring of staff.

### Treatment with bacterial surface components and cyclic lipopeptides

CoVs were diluted 10% vol/vol in solutions with final concentrations of 1 mg/ml (PG and LPS) or 100 μg/ul (CLPs) unless otherwise specified in the text. For alcohol wash experiments, samples instead diluted 5% vol/vol. Treated samples were then incubated for 2 hours at 37°C, after which they were titrated. Bacterial components were purchased from Sigma-Aldrich; lipopolysaccharides from *Escherichia coli* (L4130), peptidoglycan *Bacillus subtilis* (69554), *Staphylococcus aureus* (77140), *Streptomyces spp.* (79682), and *Micrococcus luteus* (53243). Peptidoglycan from *Escherichia coli* (PGN-EB) was purchased from Invivogen. For each surface component, stock solutions were created by suspending the component in PBS and stored and −20°C. The cyclic lipopeptides surfactin (S3523), iturin A (I1774), fengycin (SMB00292), polymyxin B (P1004), colisitn (C4461), ramoplanin (R1781), and daptomycin (D2446) were also purchased from Sigma-Aldrich.

### Mass spectrometry

Stock peptidoglycan was centrifuged at 15,000g for 1 minute in a table top centrifuge and the insoluble PG fraction was then resuspended in 100% ethanol. Following a 5-minute incubation at room temperature, samples were centrifuged, and the supernatant and insoluble fractions were used for treatment of viruses or delivered to the mass spectrometry core facility. 1 μl of peptidoglycan was combined 1:1 with a 10 mg/ml α-cyano-4-hydroxycinnamic acid (60% acetonitrile) and spotted onto MALDI targets. All MALDI-MS experiments were performed using a 5800 MALDI-TOF/TOF (Applied Biosystems). The MS data were acquired using the reflectron detector in positive mode (700–4500 Da, 1900 Da focus mass) using 300 laser shots (50 shots per sub-spectrum). Collision induced dissociation tandem MS spectra were acquired using 1 kV of collision energy. Fragmentation data was analyzed manually to determine structural information.

### Transmission Electron Microscopy

HCoV-229E virions were visualized by transmission electron microscopy (TEM) through negative staining with 2% Uranyl acetate (46). Briefly, 200 mesh formvar carbon-coated copper grids (FCF200-CU) from Electron Microscopy Sciences were treated for 20 minutes with HCoV-229E samples. Excess sample solution was then wicked off with filter paper, and each grid was then stained for 45 seconds with 2% Uranyl-acetate solution. Excess stain was again wicked off with filter paper. Grids were then dried and visualized on a Philips CM100 TEM Electron microscope. Images were recorded with a Gatan Orius SC200 CCD camera. In order to ensure even counting, 10 pictures were taken on 3 different cells on each grid. No more than 10 minutes were allotted for looking for virus in each cell.

### RNase I protection assay

Assays were performed in accordance with standard protocols described previously (47). Briefly, Samples were treated either with or without 250U RNase I for 30 minutes. To halt RNA digestion and inactivate RNase I, 2 times volume of Viral RNA Buffer from Zymo Research (R1034-1-100) with 2-mercaptoethanol was added. RNA was then extracted using the Quick-RNA Viral Kit from Zymo Research (R1035). RNA was then converted into cDNA using the iScript cDNA synthesis Kit (170-8891) from Bio-Rad. Quantitative real time PCR was performed using SsoAdvanced Universal SYBR Green Supermix (172-5271) from Bio-Rad. HCoV-229E specific primer sequences were Forward: 5-TGACATTCGCGACTACAAGC-3 and Reverse: 5-TAACGGTGGTTTGGCTTTTC-3. MERS-CoV specific primer sequences were Forward: 5-TCGCTTGGCAAATGAGTGTG-3 and Reverse: 5-ACATTAGCAGTTGTCGCCTG-3.

### Statistical analysis

All statistical comparisons in this manuscript involved the comparison between 2 groups, untreated control virus and peptidoglycan/surfactin treated virus. Thus, significant differences in viral titer, TEM counts, RNA levels, and weight loss were determined by the unpaired two-tailed students T-Test.

### Ethic Statement

This study was carried out in accordance with the recommendations for care and use of animals by the Office of Laboratory Animal Welfare, National Institutes of Health. The Institutional Animal Care and Use Committee (IACUC) of University of Texas Medical Branch (UTMB) approved the animal studies under protocol 1711065 and 1707046.

### Mice and *in vivo* infection

Ten-week old BALB/c mice were purchased from Charles River Laboratories and maintained in Sealsafe™ HEPA-filtered air in/out units. Animals were anesthetized with isoflurane and infected intranasally (IN) with 10^4^ PFU in 50 μl of phosphate-buffered saline (PBS). Infected animals were monitored for weight loss, morbidity, and clinical signs of disease, and lung titers were determined as described previously (48). For experiments l of 100 μg/ml surfactin-PBS was IN administered to anesthetized animals 18 hours prior to infection with additional treatments on day 0, day 1, and day 2. Infected animals were weighed daily, and lungs collected 2 and 4 days post-infection for downstream analysis by plaque assay.

## Acknowledgements

Research was supported by grants from NIA and NIAID of the NIH (U19AI100625 and R00AG049092 to VDM; R01AI134907, R21AI126012, R21AI132479-01 to RR; R24AI120942 to WRCEVA). Research was also supported by STARs Award provided by the University of Texas System to VDM and trainee funding provided by the McLaughlin Endowment at UTMB.

